# Accuracy of mutual predictions of plant and microbe communities varies along a successional gradient in an alpine glacier forefield

**DOI:** 10.1101/2021.08.27.457913

**Authors:** Xie He, Maximilian Hanusch, Victoria Ruiz-Hernández, Robert R. Junker

## Abstract

- Due to climate warming, recently deglaciated glacier forefields create virtually uninhabited substrates waiting for initial colonization of bacteria, fungi and plants and serve as an ideal ecosystem for studying transformations in community composition and diversity over time and the interactions between taxonomic groups.
- In this study, we investigated the composition and diversity of bacteria, and fungi, plants and environmental factors (pH, temperature, plot age and soil nutrients) along a 1.5km glacier forefield. We used random forest analysis to detect how well the composition and diversity of taxonomic groups and environmental factors can be mutually predicted.
- Community composition and diversity of taxonomic groups predicted each other more accurately than environmental factors predicted the taxonomic groups; within the taxonomic groups bacteria and fungi predicted each other best and the taxa’s composition was better predicted than diversity indices. Additionally, accuracy of prediction among taxonomic groups and environmental factors considerably varied along the successional gradient.
- Although our results are no direct indication of interactions between the taxa investigated and the environmental conditions, the accurate predictions among bacteria, fungi, and plants do provide insights into the concerted community assembly of different taxa in response to changing environments along a successional gradient.

## Introduction

Ecological successions represent a sequence of assembly processes leading to diverse and complex communities. It is widely acknowledged that in primary successions stochastic events dominate the initial community assembly whereas niche-based processes become more important in further developed communities where both the environment and species interactions shape species composition and diversity (Chang & HilleRisLambers, 2016; Wojcik *et al*., 2021). Climate warming is initiating primary successions in glacier forefields as it is speeding up glacier retreat and providing newly uninhabited substrates for the colonization of organisms forming communities along the chronosequence. Local microclimatic conditions and soil properties, and the tight interactions between plant and belowground microbes are part of the environmental and biotic factors shaping these communities (Zak *et al*., 2003; Mouhamadou *et al*., 2013; Darcy *et al*., 2018; Navratilova *et al*., 2019; Harrison *et al*., 2020; Ohler *et al*., 2020; Davison *et al*., 2021). The relative contributions of environmental and biotic factors on communities may vary spatially and temporally and may affect different properties of communities (Mitchell *et al*., 2011; Chen *et al*., 2017). In the case of microbial communities the relative importance of these factors partly depends on the soil compartment: Microbes colonizing the rhizosphere are directly affected by the sugars, organic acids and amino acids provided by root exudates, whereas microbes colonizing bulk soil are less affected by specific plant individuals and may thus respond more strongly to the environmental conditions (Hartley *et al*., 2007; Lange *et al*., 2015; Sanchez-Canizares *et al*., 2017). Accordingly, both environmental factors such as soil chemical properties and temperature (Cheng *et al*., 2020; Hermans *et al*., 2020; Davison *et al*., 2021) as well as plant community composition (Chen *et al*., 2017; Kruger *et al*., 2017; Reese *et al*., 2018) were reported to explain variation in soil microbial communities. Likewise, microbes and the environment have been shown to affect plant species composition (Bever *et al*., 2012; Miller *et al*., 2020). For instance, recent studies demonstrated that bacterial and fungal communities in soil may either positively or negatively affect plant species and communities, and that these effects are tightly related to the plant functional type or the partnership between the plants and microbial organisms (Teste *et al*., 2017; Hahl *et al*., 2020; Heinen *et al*., 2020). Within soil microbial communities, the various interactions between bacteria and fungi are additionally contributing to community assembly. The interactions between microbes that share a habitat can either be positive (mutualism, synergism, or commensalism), negative (pathogenic, predation, parasitism, antagonism, or competition) or neutral (no observed effects) (Vandenkoornhuyse *et al*., 2007; Berg *et al*., 2020). In diverse communities all of these outcomes of pairwise interactions may occur simultaneously, which lead to changes of the organism performance and ecosystem productivity (Wargo & Hogan, 2006; Miransari, 2011). The outcome of pairwise interactions between plant, bacterial, and fungal species is highly context dependent and thus modulated by the presence of other species as well as environmental conditions. For instance, environmental conditions like temperature and soil moisture affect plant and microbes and can regulate plant-microbe associations (Rasmussen *et al*., 2019; Rudgers *et al*., 2020; Robroek *et al*., 2021), and increasing environmental stress alters microbial facilitation of plant germination or biomass production (David *et al*., 2020).

Successional gradients such as glacier forefields with considerable variation in soil properties and climate conditions are an ideal study system to reveal how the interdependences between taxonomic groups change with environmental conditions. Previous studies have shown different interactive patterns of organisms in different successional stages. For instance, a positive relationship between plant and microbial richness was found in the early succession while not in the late succession (Porazinska *et al*., 2018), which may be owing to the fact that pioneering plants serve as nutrition hotspot for microbes in the rhizosphere at the initial sites (Schulz *et al*., 2013) while in the late succession the accumulated organic matter provide plenty resources and thus the mutual predictability between plants and microbes is reduced (Porazinska *et al*., 2018). Nonetheless, the interdependencies between the organisms that form complex communities as well as environmental conditions may also leave a signal in community composition and diversity of plants and microbes and thus these properties may be mutually predictable (Horn *et al*., 2017; Leff *et al*., 2018).

Machine learning algorithms have been increasingly applied for pattern recognition and predictions using complex ecological data. For instance, random forest analysis was used to explore the links between soil bacterial community composition and environmental factors such as land use management and soil properties and lead to predictive models with high accuracy (Hermans *et al*., 2020). Furthermore, machine learning models were shown to outperform regression models in trait-matching predictions for understanding interaction networks (Pichler *et al*., 2019). The high performance of machine learning algorithms and especially random forest is obtained by their ability to model non-linear combinations of numerical and categorial data without complex transformations resulting in estimates of the accuracy of predictions as well as the importance of individual variables in improving predictions (Breiman, 2001; Ghannam & Techtmann, 2021). Thus, random forest is an excellent tool for evaluating the interdependences between various taxonomic groups and environmental factors (Ghannam & Techtmann, 2021; Goodswen *et al*., 2021).

In recent studies, individual plant variables such as species composition, functional identity, taxonomic, phylogenetic and functional diversity have been used to predict microbial communities (Prober *et al*., 2015; Dassen *et al*., 2017; Chen *et al*., 2018; Leff *et al*., 2018; Porazinska *et al*., 2018). In the present study, we consider all the possible variables and evaluate how well the diversity and composition of plants, bacteria, fungi, and environmental factors can predict each other in order to explore the interdependences between the taxonomic groups and environmental factors as well as their changes along a successional gradient in the forefield of the Ödenwinkelkees glacier in the Austrian Alps (Junker *et al*., 2020). Here the assembly of multidiverse communities along the glacier forefield chronosequence provides an excellent opportunity to track transformations in community composition and diversity over time. Using random forest analysis, we used plant, bacteria, fungi composition and environmental factors as explanatory variables and bacteria, fungi and plant species composition, plant functional composition, plants phylogenetic, functional diversities and environmental factors as dependent variables to test for interdependencies between these variables. We aim to address the following two questions: 1) How accurately can we predict properties of plant and microbial communities with the composition of the other taxonomic groups as well as environmental factors? 2) Is the accuracy of prediction variable along the successional gradient? Although our approach does not directly indicate interactions and dependencies between taxonomic groups and environmental conditions, it tests the hypothesis that taxonomic groups respond to changing environmental conditions in a concerted way potentially facilitated by tight interaction networks.

## Materials and methods

### Data collection

Our study site was located at the forefield of the Ödenwinkelkees glacier (Stubachtal valley, Hohe Tauern National Park, Austria; Dynamic Ecological Information Management System – site and dataset registry: https://deims.org/activity/fefd07db-2f16-46eb-8883-f10fbc9d13a3, last access: August 2021) (Junker *et al*., 2020). In summer 2019 (26 June - 16 September), we established 135 permanent plots within the successional gradient of the glacier forefield. We identified all vascular plant species occurring at the plots (*n* = 107) and recorded the coverage of plants with a resolution of 0.1%. We measured the plant height, leaf area, leaf weight and calculated the specific leaf area (SLA) for those 48 plant species that occurred in 10 or more plots. For three focus species we phenotyped up to three individuals on every plot where they occurred: *Oxyria digyna* as representative of early succession, *Trifolium badium* as representative of late succession, and *Campanula scheuchzeri* which occurred all along the successional gradient (for detailed information on the selection of the focus plant species see Junker et. al 2020). For the other *n* = 45 species, up to five individuals per plot were phenotyped on the youngest, the oldest, and the intermediate plot where they occurred (for detailed methods see Junker et al 2020). Additionally, we obtained the functional traits of the plant species from Bioflor database (https://www.ufz.de/biolflor/index.jsp) for 92 species out of 107 plant species occurring in the field. We used nine functional traits which have been shown to be response traits to environmental changes at the community level (Kahmen & Poschlod, 2004; Bernhardt-Römermann *et al*., 2008; Aguiar *et al*., 2013; Hintze *et al*., 2013), including fruit type, leaf anatomy, leaf persistence, life form, life span, pollen vector, strategy type, type of reproduction. We also characterized the soil microbiome (bacteria and fungi) of each of the plots. We sampled soil from each plot at two locations at a depth of 3cm, soil from two locations per plot were pooled to one sample for further analysis. Samples were directly transferred to ZR BashingBeads Lysis tubes containing 750 µL of ZymoBIOMICS lysis solution (Zymo-BIOMICS DNA Miniprep Kit; Zymo Research, Irvine, California, USA). Within 8h after collection of microbial samples, ZR BashingBeads Lysis tubes were sonicated for 7 min to detach microorganisms from the surfaces. Subsequently, all microbial samples were shaken using a ball mill for 9 minutes with a frequency of 30.0 s^−1^. Microbial DNA was extracted using the ZymoBIOMICS DNA Miniprep Kit following the manufacturer’s instructions. Microbiome analysis was performed by Eurofins Genomics (Ebersberg, Germany) using the company’s standard procedure. To assign taxonomic information to each OTU, DC-MEGABLAST alignments of cluster representative sequences to the sequence database were performed (Reference database: NCBI_nt (Release 2018-07-07)). Further processing of OTUs and taxonomic assignments was performed using the QIIME software package (version 1.9.1, http://qiime.org/) (Caporaso *et al*., 2010). Abundances of bacterial and fungal taxonomic units were normalized using lineage-specific copy numbers of the relevant marker genes to improve estimates (Angly *et al*., 2014). Prior to the statistical analysis of microbial communities, we performed a cumulative sum scaling (CSS) normalization (R package metagenomeSeq v1.28.2) on the count data to account for differences in sequencing depth among samples.

To record the seasonal mean temperature, we buried temperature loggers with a resolution of 0.5 °C (MF1921G iButton, Fuchs Elektronik, Weinheim, Germany) 10 cm north of each plot center, at a depth of 3 cm below ground (Junker *et al*., 2020; Ohler *et al*., 2020) during field work in 2019. The thermo loggers were set to start on 13th August 2019 and were stopped on 9th August 2020 with a total of 2048 measurements recorded on 362 days. Seasonal mean temperature was calculated on the basis of the recordings ranging from 13th August to 16th of September 2019 and 26th June to 9th August 2020 representing the period in which the plots were free of permanent snow cover before and after the winter 2019/2020. In 2020 (25 July - 21 August), we took additional soil samples from all plots to measure soil nutrient content (N P, K, Mg) as well as soil pH. Samples were sent to AGROLAB Agrar und Umwelt GmbH (Sarstedt, Germany) for analysis.

### Data analysis

To test the predictability of the diversity and composition of each of the taxonomic group by the composition of other taxonomic groups as well as by environmental parameters, we used the machine learning algorithm random forest (R package randomForest). Random forest combines several randomized decision trees and aggregates their predictions by averaging, it can handle multiple input variables (explanatory variables), which are ranked by different levels of importance in predicting the dependent variable (Breiman, 2001; Biau & Scornet, 2016). As explanatory variables we used the community tables of plants, bacteria, and fungi with plots as rows and the abundance of the species or OTUs as columns (Table S1, S2 & S3); meanwhile we used multivariate datasets informing about the environmental conditions of each plot with plots as rows and environmental variables as columns (Table S4). As dependent variables we used univariate variable including plant Shannon, phylogenetic and functional diversity, bacteria Shannon diversity, fungi Shannon diversity, soil seasonal mean temperature, pH, plot age, soil N, P, K, and Mg as well as principal components of the composition of all the taxonomic groups, resulting in 20 variables in total (Table S5). As random forest analysis can only deal with univariate dependent variables, we used the first two principal component axis (PCA) which carry most information of the composition to refer to plant species composition (15.3% + 11.2%), bacteria composition (6.4% + 4.6%) and fungi composition (4.1% + 3.2%). Plant functional composition matrix was generated based on the plant species composition table and the functional traits table obtained from Bioflor database. For each category of each trait, we calculated the total coverage of species belonging to the category, and this was done for all the 9 traits and all 9 traits were merged to a single table, thus generating the functional composition table with plots name as rows and 50 trait categories as columns, i.e. each categorial traits had two or more categories resulting in a total of 50 categories. (Table S6). Plant functional composition was represented by the first two PCAs, too (59.3% + 12.3%). Plant Shannon diversity was calculated from the compositional dataset using the R package vegan (Dixon, 2003). Plant phylogenetic diversity was calculated using the R package picante (Kembel *et al*., 2010). We extracted a phylogenetic tree using the R package pez (Pearse *et al*., 2015) for species existing in our field site from a dated molecular phylogeny tree (32,223 species) for land plants (Zanne *et al*., 2014). In cases where species were not included in the tree, it was substituted by species from the same genus. Among 107 species existing in our plots, we were able to match and built a tree with 104 species and we used it for the calculation of phylogenetic diversity. We used ‘Functional dispersion’ calculated from the R package FD (Laliberte & Legendre, 2010) as the index for plant functional diversity. The mean plant height, leaf area, leaf weight and SLA of every species were used for the trait table (Table S7) identically for every plot, and for the community table the species with a low occurring frequency along the successional gradient (not included in the 48 species with traits measured) were ignored in the calculation of functional diversity. For bacteria and fungi, the Shannon diversity was calculated based on the OTU composition after rarefying the data to the minimum number of reads available in the samples (repeats = 999).

Using all combinations of explanatory and dependent variables, we performed random forest analyses with 10-fold cross validations to quantify the performance of the predictive model. Specifically, for each prediction, 80% of the plots were randomly selected as the training dataset and the remaining 20% of the plots were used as test dataset. The predictive model resulting from the training dataset was applied to the test data and the predicted values of the plots in the test dataset were correlated with the observed values of these plots. This process was repeated for ten times, and then we defined the mean Pearson’s *r*-value of ten correlations as ‘accuracy of prediction’ and used the proportion of significant correlations (p-value < 0.05) out of the 10 correlations as ‘significance frequency’. Additional to random forest analysis using all the plots for a global impression on the predictability of dependent variables, we also employed a moving frame approach to detect how the predictabilities change along the successional gradient. With the 135 plots, we grouped every 45 plots into one frame and used the median plot as identifier of the frame. Thus, the first frame included plots 1 to 45, the second 2 to 46, and so forth. This approach led to a set of 91 moving frames whose identifiers ranged from plot 23 to plot 113. Using the same proportion of training and test dataset, for every 45 plots in each frame, data of 36 (80%) randomly selected plots was used as training dataset, and the other 9 (20%) plots were used as test dataset. The accuracy of prediction and significance frequency were calculated for every frame as stated before. We fitted a linear or quadratic regression with the accuracy of prediction of every variable along the successional gradient as independent variable and the frame number as explanatory variable. The model with a higher *r*^*2*^ value was chosen and the statistically significant relationships were shown as a regression line.

To make a comprehensive comparison of how well every variable is predicted by the other individual group and by the other three groups combined, we did the same random forest predicting procedure for each of the 20 variables using the other three groups together (except for the group that was considered in the dependent variable). We compared for each variable how well they were predicted by every other single group and by three groups combined using the Tukey Test.

## Results

In total we obtained soil bacteria and fungi composition data from *n* = 127 and 130 plots, respectively; *n* = 5221 bacteria OTUs and *n* = 6016 fungi OTUs were detected in all the soil samples. Raw sequences of next-generation 16S rRNA gene amplicon sequencing are available at the NCBI Sequence Read Archive (SRA) under the BioProject accession PRJNA701884 and PRJNA701890. The mean accuracy of prediction of each pair of explanatory variables and dependent variables did usually not strongly differ between the global analysis considering all plots and the mean of the frame-wise analyses, indicating the validity of using the moving frames for random forest predictions. Most of the predictions fit a quadratic regression, indicating a non-monotonic change of the accuracy of prediction along the successional gradient.

### Bacterial communities as predictors (Fig. 1 and Fig. 5a)

Bacterial communities (quantitative OTU tables) most accurately predicted the taxonomic composition of fungal communities (PC1 and PC2) followed by plant functional composition. Among the environmental parameters, plot age and pH-value were most accurately predicted by bacterial communities. Note that our results do not imply a direction of effects in the sense that the dependent variable is affected by the explanatory variable. For instance, bacterial communities do not affect the soil temperature but are affected by this environmental parameter. Accuracy of prediction of target variables associated with plant communities mostly decreased with plot age, whereas accuracy of prediction of fungi and environmental target variables remained constant or even increased along the age gradient in most cases.

**Fig. 1.**
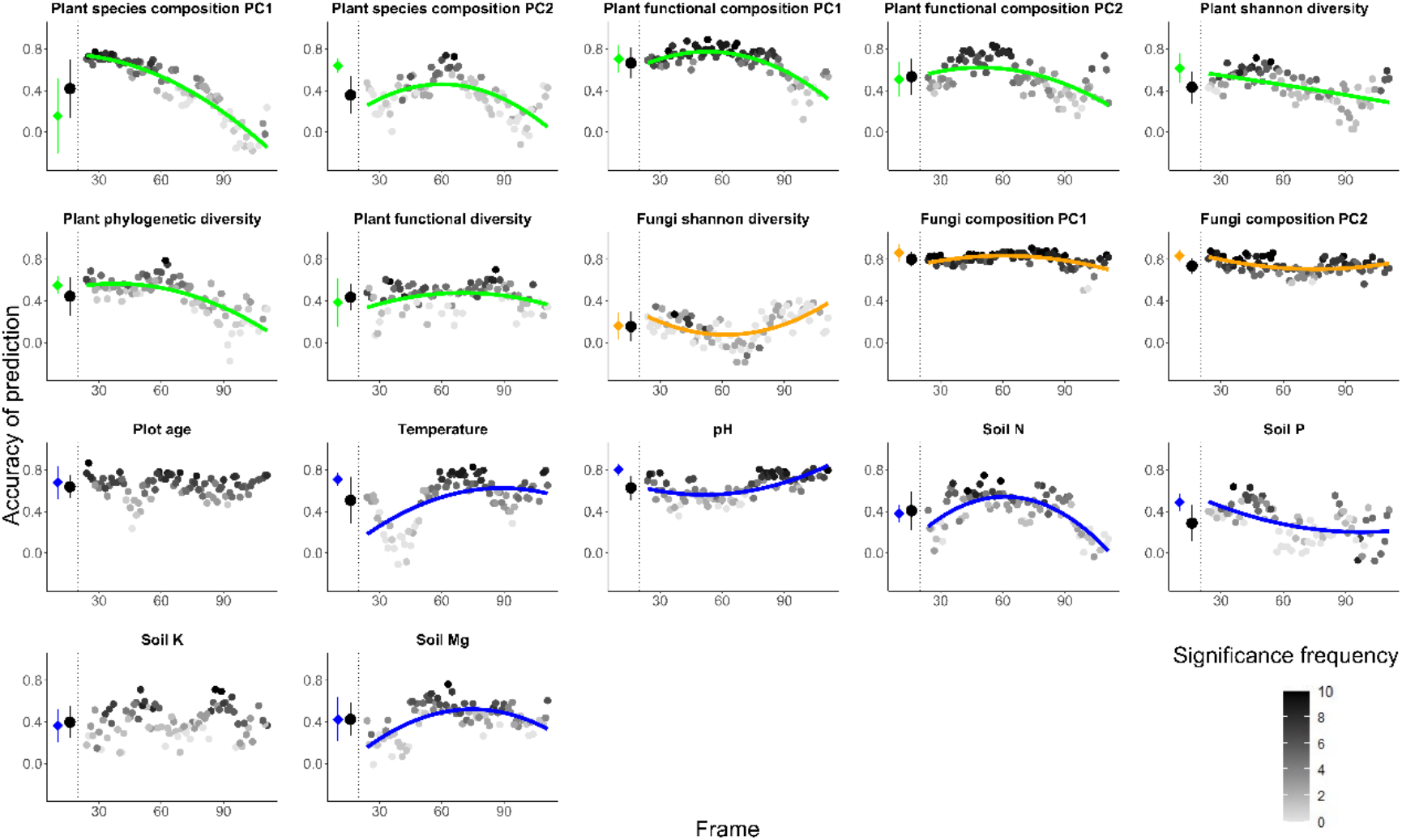
Random forest predictions using the community table of soil bacterial communities (OTU table) to predict seven variables of plant (green) and three variables of fungi (orange) as well as seven variables of environmental factors (blue). The colored circles at the left of each plot denote the mean ± standard deviation of the accuracy of prediction using the full dataset (results of 10-fold cross validation), and the black circles denote the mean ± standard deviation of the accuracy of prediction for all the frames. Each grey to black circle on the right of each plot represents the mean accuracy of prediction of each frame and the color gradient is showing how many correlations of the 10-fold cross-validation were significant with lighter colors indicating less frequent significant predictions. A quadratic or linear regression (the model with higher adjusted r2 value) is fit for the gradient if it is significant, showing a change of the accuracy of prediction along the successional gradient.

### Fungi communities as predictors (Fig. 2 and Fig. 5b)

Fungal communities (quantitative OTU tables) most accurately predicted the taxonomic composition of bacterial communities (PC1 and PC2) and bacterial Shannon diversity was the variable with the lowest accuracy of prediction. Plot age and pH were also the environmental factors that were most accurately predicted by fungi communities. Similar to bacterial predictions, accuracy of prediction of target variables associated with plant communities mostly decreased with plot age, whereas accuracy of prediction of bacterial and environmental target variables remained constant or increased along the age gradient in most cases.

**Fig. 2.**
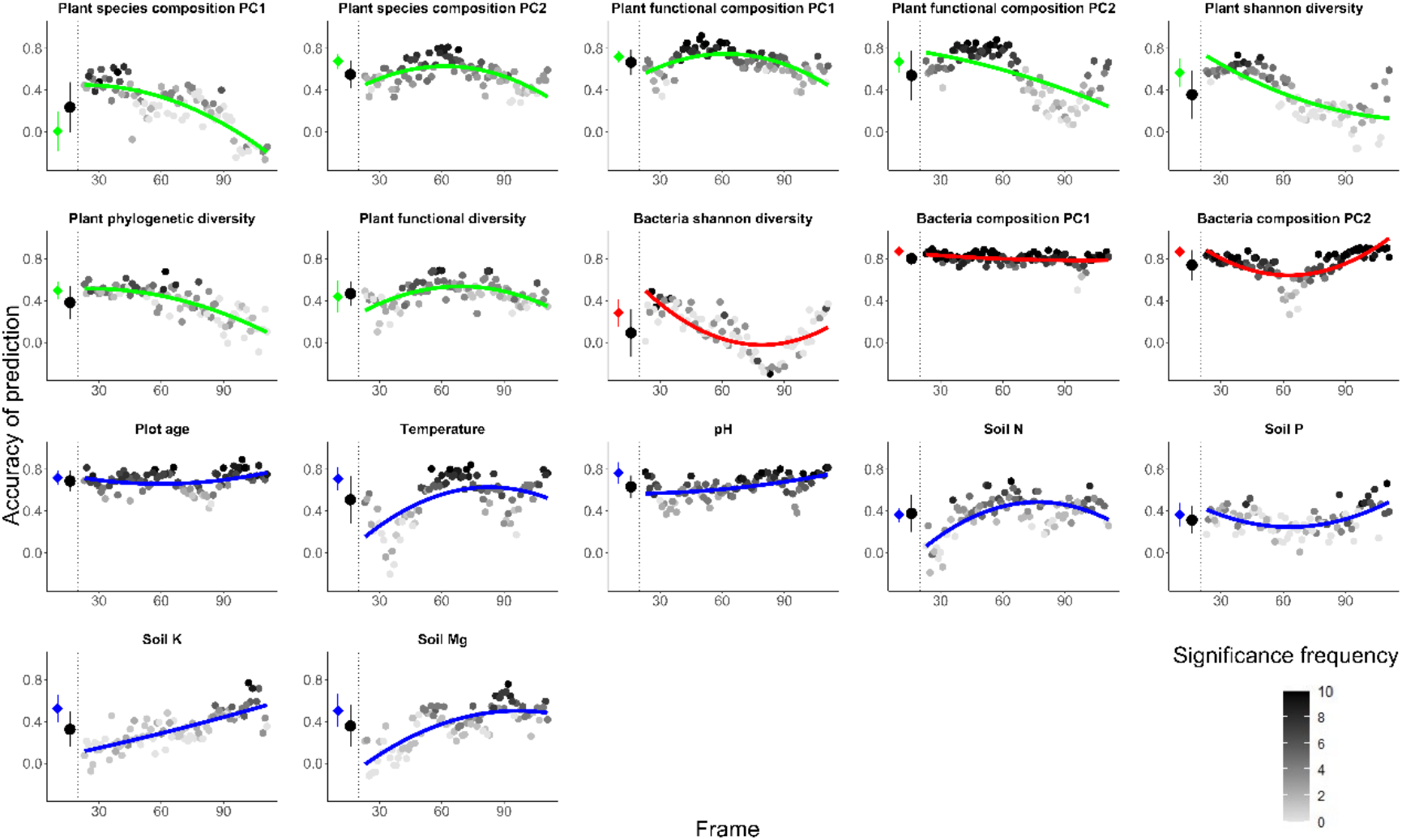
Random forest predictions using the community table of soil fungal communities (OUT table) to predict seven variables of plant (green) and three variables of bacteria (red) as well as seven variables of environmental factors (blue). The colored circles at the left of each plot denote the mean ± standard deviation of the accuracy of prediction using the full dataset (results of 10-fold cross validation), and the black circles denote the mean ± standard deviation of the accuracy of prediction for all the frames. Each grey to black circle on the right of each plot represents the mean accuracy of prediction of each frame and the color gradient is showing how many correlations of the 10-fold cross-validation were significant with lighter colors indicating less frequent significant predictions. A quadratic or linear regression (the model with higher adjusted r2 value) is fit for the gradient if it is significant, showing a change of the accuracy of prediction along the successional gradient.

### Plant communities as predictors (Fig. 3 and Fig. 5c)

Plant communities (quantitative vegetation table) predicted the plot age most accurately, followed by fungi composition (PC1) and bacteria composition (PC1). Plant communities predicted bacteria and fungi Shannon diversities least accurately. The plant predictions of variables concerning bacteria, fungi and some environmental parameters were mostly decreasing with increasing plot age. For environmental variables, the accuracy of prediction for temperature, pH and soil Mg increased and the others were mostly decreasing with plot age.

**Fig. 3.**
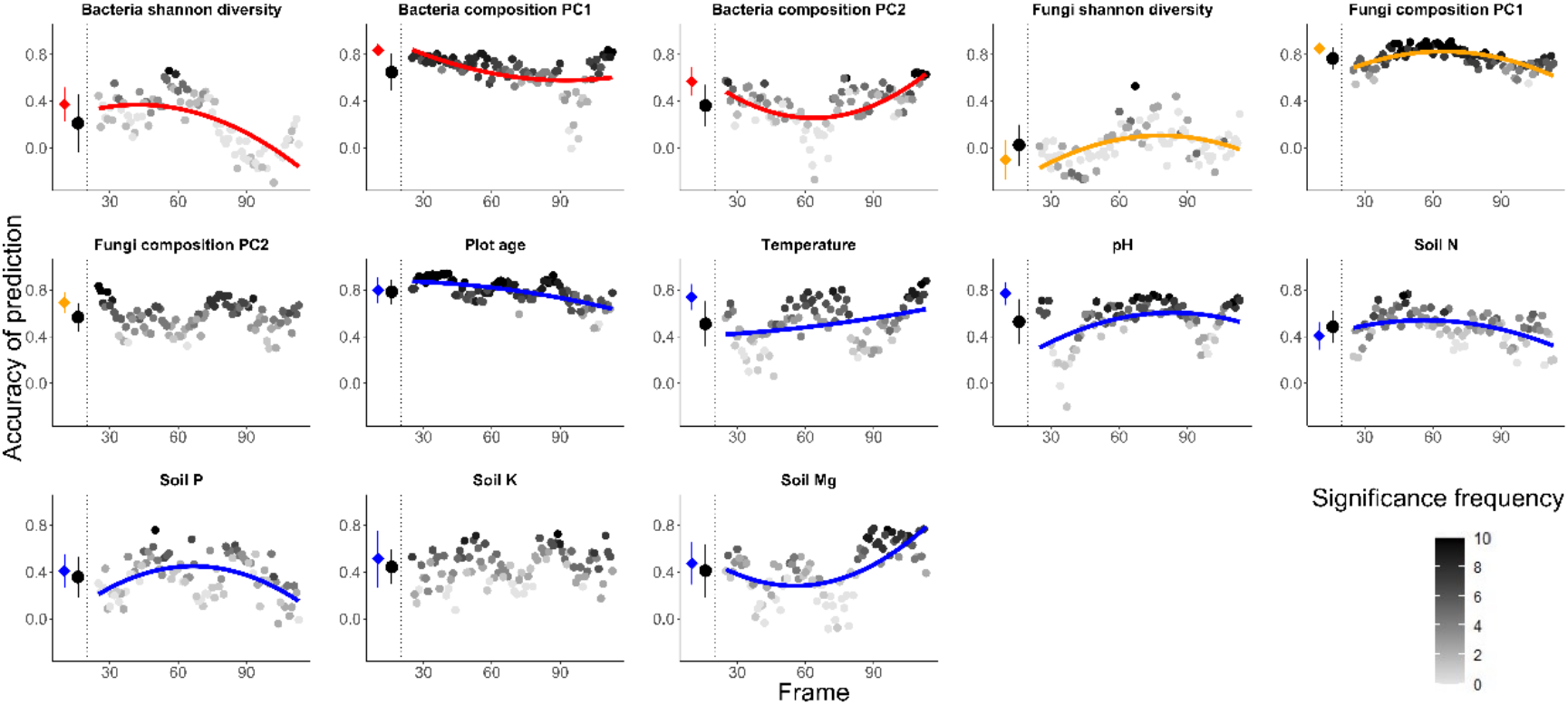
Random forest predictions using the community table of plant communities to predict three variables of bacteria (red) and three variables of fungi (orange) as well as seven variables of environmental factors (blue). The colored circles at the left of each plot denote the mean ± standard deviation of the accuracy of prediction using the full dataset (results of 10-fold cross validation), and the black circles denote the mean ± standard deviation of the accuracy of prediction for all the frames. Each grey to black circle on the right of each plot represents the mean accuracy of prediction of each frame and the color gradient is showing how many correlations of the 10-fold cross-validation were significant with lighter colors indicating less frequent significant predictions. A quadratic or linear regression (the model with higher adjusted r2 value) is fit for the gradient if it is significant, showing a change of the accuracy of prediction along the successional gradient.

### Environmental factors as predictors (Fig. 4 and Fig. 5d)

Environmental factors (multivariate table of environmental parameters) predicted the fungi composition PC1 and bacteria composition PC1 with the highest accuracy, followed by plant functional diversity and plant species composition PC2. Accuracy of prediction for plant variables were mostly decreasing along the gradient, and for bacteria and fungi they either had the highest accuracy of prediction in the middle age or increase with plot age.

**Fig. 4.**
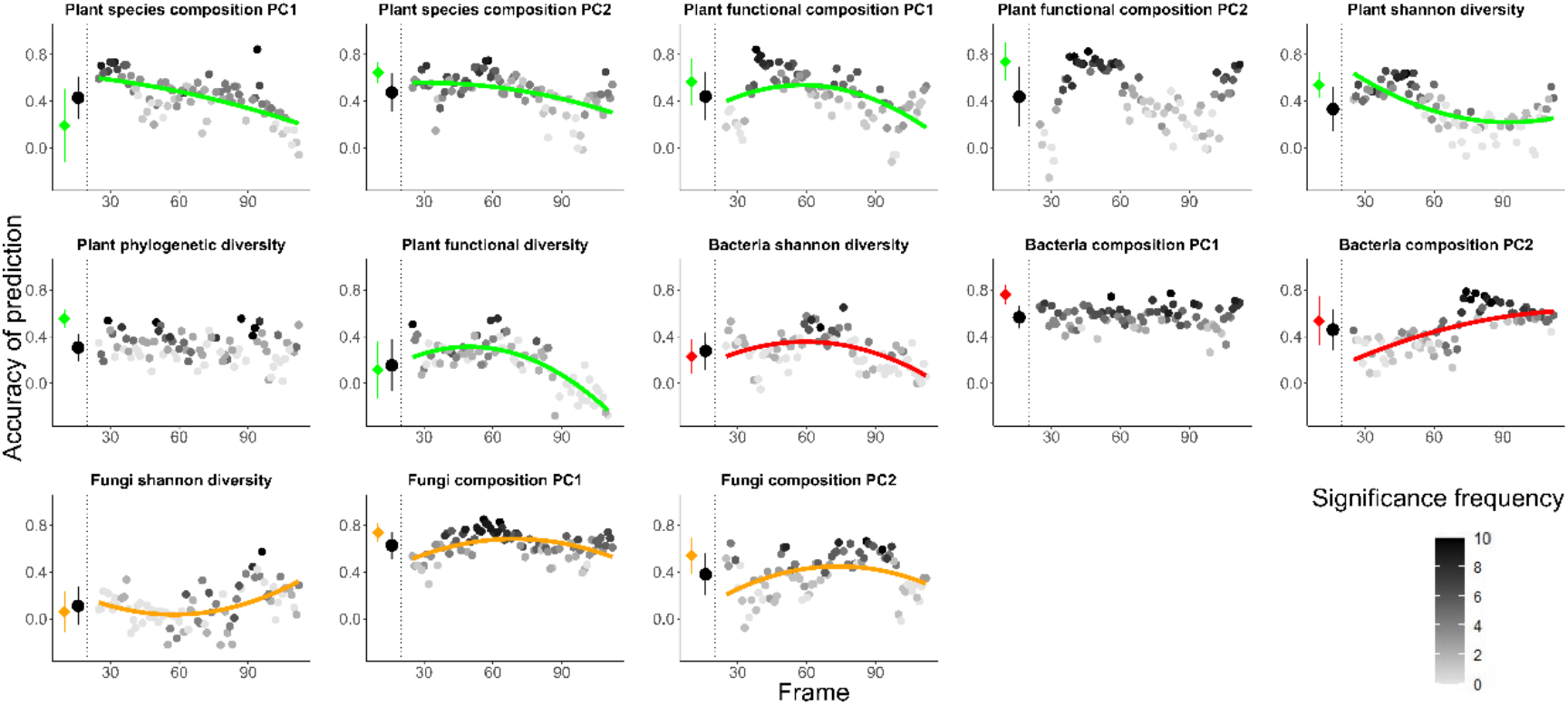
Random forest predictions using all the environmental factors to predict seven variables of plant (green) and three variables of bacteria (red) as well as three variables of fungi (orange). The colored circles at the left of each plot denote the mean ± standard deviation of the accuracy of prediction using the full dataset (results of 10-fold cross validation), and the black circles denote the mean ± standard deviation of the accuracy of prediction for all the frames. Each grey to black circle on the right of each plot represents the mean accuracy of prediction of each frame and the color gradient is showing how many correlations of the 10-fold cross-validation were significant with lighter colors indicating less frequent significant predictions. A quadratic or linear regression (the model with higher adjusted r2 value) is fit for the gradient if it is significant, showing a change of the accuracy of prediction along the successional gradient.

**Fig. 5.**
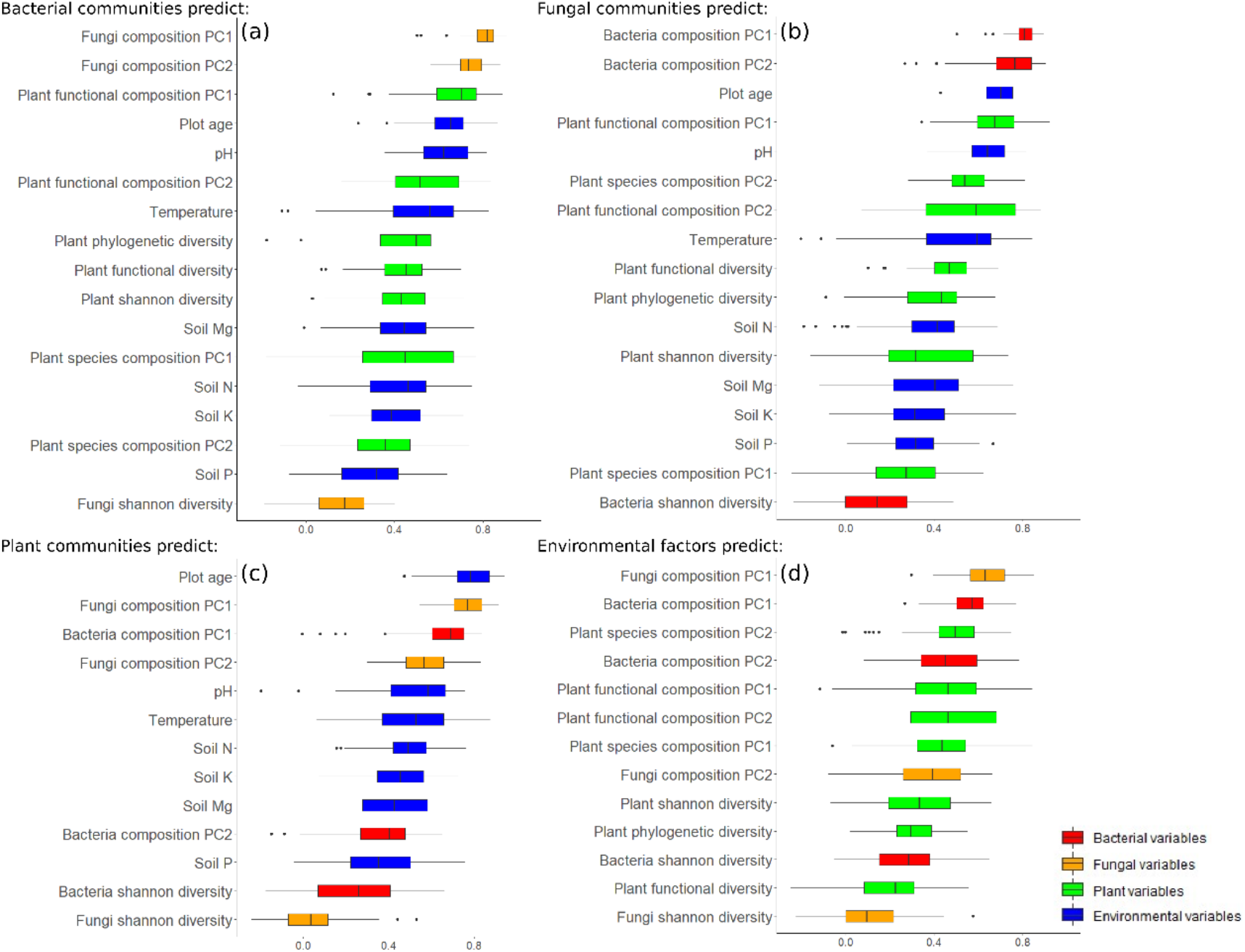
Summary of the accuracy of prediction using taxonomic groups (bacteria (a), fungi (b), plant (c)) and environmental factors (d) to predict variables from the other three groups along the successional gradient. Variables from each group are color-coded (red: bacteria, orange: fungi, green: plant, blue: environment) and ranked by accuracy of prediction.

### Combined groups as predictors (Fig. 6)

Using all taxonomic groups and environmental variables (except for the group that was considered in the dependent variable) together to predict dependent variables increased the accuracy of prediction for plant functional composition PC1, plant functional composition PC2, fungi composition PC1. For plot age, the accuracy of prediction even decreased, and for other variables especially environmental variables, the accuracy of prediction with all the groups combined did not significantly increase.

**Fig. 6.**
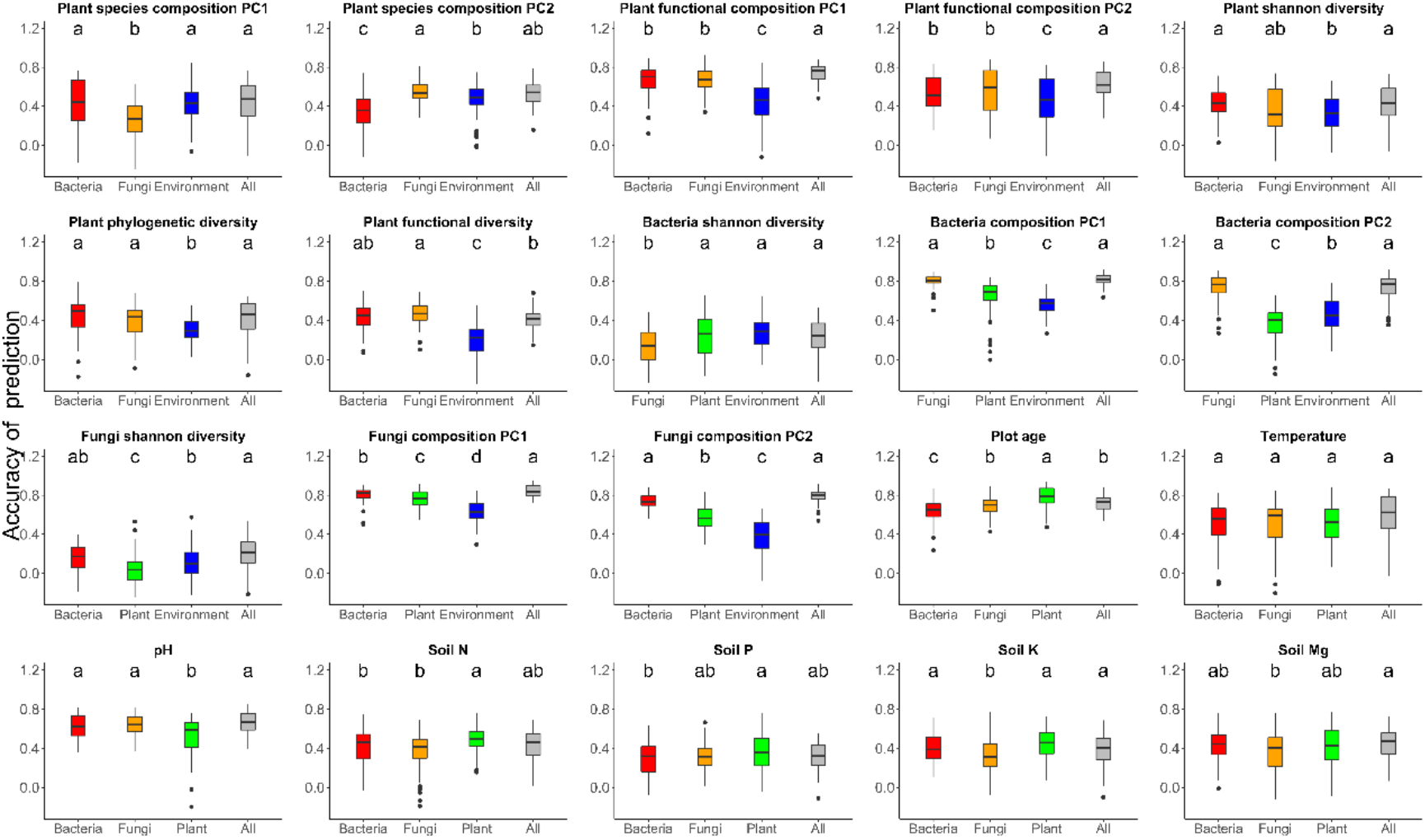
Summary of the accuracy of prediction for each variable being predicted by every single group (red: bacteria, orange: fungi, green: plant, blue: environment) as well as by the other three groups combined (grey). The label on each boxplot is the result of Tukey Test showing if there is significant difference of accuracy of prediction between any pair of predicting groups.

## Discussion

Our results indicate that the composition and diversity of plant, bacteria, and fungi is - to a certain degree - predictable by the composition of the respective other taxonomic groups as well as by environmental factors. The accuracy of prediction, however, varied along the successional gradient of the forefield of the Ödenwinkelkees glacier. Overall, the taxonomic groups predicted each other more accurately than environmental factors predicted the taxonomic groups; within the taxonomic groups their composition was better predicted than diversity indices. When using a combined dataset as predictors, only a few variables obtained increased accuracy of prediction compared with using a single group, and most of the variables have no significant difference or even decreased accuracy of prediction. Well performing predictive models may indicate direct interactions between taxa or effects of the environment on taxa. However, statistical associations between taxa may also suggest that both taxa respond similarly to a third taxonomic group or an environmental factor (Blanchet *et al*., 2020). Thus, while our results do not directly reveal ecological interactions, they do provide insights into the concerted community assembly of different taxa in response to changing environmental factors along a successional gradient.

Variables describing the composition of taxonomic groups (e.g. PC axis of community composition) were mostly more precisely predicted by other taxonomic groups than diversity indices. Particularly, the community composition of bacteria and fungi mutually predicted each other most precisely, which confirms previous studies demonstrating the interdependences between bacteria and fungi (Miransari, 2011; Deveau *et al*., 2018). Both bacteria and fungi community composition predicted plant functional composition more precisely than species composition and functional, phylogenetic and taxonomic diversity of plants. These results indicate that the plant functional identity has a stronger effect on soil microbial communities than plant species identity and diversities (Dassen *et al*., 2017). Fungal composition was better predicted by plant composition than bacterial composition, which may reflect the tight interaction between plants and fungi, especially mycorrhiza (Horn *et al*., 2017; Sweeney *et al*., 2021). The interactions between plants and microbes are mediated through plant root exudates and litter input (Knelman *et al*., 2012; Lopez-Angulo *et al*., 2020). Root exudates vary substantially between different plant species and various microbes utilize the carbon source from plants (Vandenkoornhuyse *et al*., 2007). In this way, the plant community provides various niches for the microbes and plays an important role shaping microbial communities in the soil (Bever *et al*., 2012). Likewise, the interplay of facilitative and antagonistic effects determines the direction of feedbacks from soil microbes to plants and maintains the diversity of plant communities (Bever *et al*., 2012; Teste *et al*., 2017; Mony *et al*., 2021). Nevertheless, although it has been reported that plant composition has an effect on microbial richness (Lopez-Angulo *et al*., 2020), we did not detect a strong accuracy of prediction between plant composition and bacterial or fungal Shannon diversity. This suggests that interactions within taxonomic groups are reducing the accuracy of prediction between the composition and diversity of plants and microbes. For instance, positive or negative effects of individual bacterial strains on plant growth may be changed by the presence of other strains (Raza *et al*., 2020), which may lead to a hardly predictable complexity of interdependencies and influences.

Plot age, soil temperature and soil pH were well predicted by taxonomic groups, and soil nutrients were less well predicted. In contrast, the environmental variables did not accurately predict the composition and diversity of the taxa. As stated above, our approach is not implying a direction of effects, which means that it is more likely that the environmental factors affect the composition and diversity of the taxonomic groups and not *vice versa*. Among all the environmental factors, plot age is the environmental factor best predicted by taxonomic groups, followed by soil temperature and pH. Plants predicted plot age better than bacteria and fungi, and the signal was even blurred when using all the groups together. This indicates that plant communities follow a clear succession with age-specific stages. Microbes may be more responsive to other environmental factors that may act on short term fluctuations such as temperature, which is equally well predicted by the compositions of bacteria, fungi and plants suggesting its common importance in defining the niche of all taxa. Previous studies demonstrated that plants and microbes from different origins may respond to increased temperature variously, thus we may infer that climate change will shift the interactive patterns between species (Rasmussen *et al*., 2019; Rudgers *et al*., 2020; Losapio *et al*., 2021). In addition, pH was better predicted by bacteria and fungi than by plants, indicating that pH is affecting soil microbes more than plants, which is in agreement with previous studies illustrating the importance of pH in affecting microbial communities (Knelman *et al*., 2012; Shen *et al*., 2020). Soil nutrients such as N, P, K, Mg were more accurately predicted by plants than by microbes suggesting strong feedbacks between soil nutrients and plant communities (Fischer *et al*., 2019). In summary, we showed that plants, bacteria, and fungi mutually predict each other’s diversity and community composition and that environmental parameters are also well-suited predictors for the same biotic dependent variables. This is in line with previous studies demonstrating that plant communities and environmental factors are contributing and explaining different parts of variation in soil microbial communities (Mitchell *et al*., 2011) and that interactions between plants and microbes can be independent on environmental changes (Sweeney *et al*., 2021).

Accuracy of prediction varied with successional age. For instance, plant taxonomic and functional composition was better predicted by bacteria and fungi at early than late succession. This could be explained by a relatively clear signal of interaction between individual plant species and microbes at early succession while the signal of individual plant species may be diluted at late successional stages where communities become complex (Porazinska *et al*., 2018). The interactions between plants and microbes are known to be responding to primary successions. For instance, while plant-derived carbon becomes a major source for bacteria after 50 years of succession, these communities utilize ancient carbon in the first decades after deglaciation, which has been demonstrated in the area of our study site (Bardgett *et al*., 2007). In accordance with this finding, Tscherko *et al*. (2005) found evidence of plants shaping microbial communities in soils older than 43 years in another Austria glacier, the Rotmoosferner. These results suggest a higher accuracy of prediction of microbial communities by plants at later successional stages, which is not fully in line with our findings. In contrast to many other statistical methods, random forest decision trees consider individual features instead of multivariate representations of the communities. Thus, even though bacterial communities are mainly shaped by abiotic factors and non-plant related carbon sources, random forest is able to select those strains that may be associated with the few plant species colonizing the young plots, which represents a strong signal in the data. In contrast in older plots, when plants provide the major carbon source, the signal of each individual species may be diluted resulting in a poor prediction. Additionally, further carbon sources accumulate such decomposed soil organic matters, which again sustains microbial communities unrelated to plant species diversity and composition. Another reason for our finding may be the reduced variability in plant species composition and diversity between older plots, which is a common finding in primary successions (Ortiz-Alvarez *et al*., 2018). This could also partly explain the decrease of accuracy of prediction between plants and microbes along the succession as the decreased variation of community composition makes it less sensitive to detect the change of the interacting taxa. Finally, age is not the only factor that is affecting the successional age of plots in glacier forefields, instead allogenic factors may reset successions or at least slow down successional progress in community development (Wojcik et al. 2021). These allogenic factors, such as geomorphic events, accumulate over time and thus may lead to outliers in community composition. If these outlier plots are part of test dataset, they cannot be predicted on models as predictions are only possible in the range of the training dataset.

Our results demonstrate the concerted development of plants and microbial communities regulated by environmental factors along a successional gradient, which suggests strong interdependencies between the taxa. As a next step, approaches like the one described here may be used to identify indicator species and environmental variables that inform best about the diversity and composition of ecosystems, which facilitates monitoring and conservation efforts. Additionally, climate warming demands the prediction of ecosystem-wide responses and our data presents existing patterns and offers information for future predictions.

## Acknowledgement

We thank the Hohe Tauern National Park Salzburg Administration and the Rudolfshütte for organizational and logistic support, the governing authority Land Salzburg for the permit to conduct our research (permit # 20507-96/45/7-2019). Jan-Christoph Otto, Tobias Seifert, and Anna Vojtkó for help in the field. Hamed Azarbad, Lisa-Maria Ohler and Verena Zieschank provided valuable comments to improve the study. The study was supported by the START-program of the Austrian Science Fund (FWF) granted to Robert R. Junker (Y1102).

## Author contributions

RRJ conceived the study. XH, MH, VRH and RRJ designed the study and collected the data. XH and RRJ analyzed the data. XH and RRJ wrote the manuscript with critical input from MH and VRH. All authors contributed to the manuscript and approved the final version.

## Supporting Information

Additional Supporting Information may be found online in the Supporting Information section at the end of the article.

**Table S1** Species composition of plant communities used as explanatory variables in predictive models.

**Table S2** Community table of soil bacterial communities (OTU table) used as explanatory variables in predictive models.

**Table S3** Community table of soil fungal communities (OTU table) used as explanatory variables in predictive models.

**Table S4** Environmental parameters table used as explanatory variables in predictive models.

**Table S5** All the 20 dependent variables used for predictive models.

**Table S6** Functional composition of plant communities.

**Table S7** Normalized mean value of field-measured traits (plant height, leaf area, leaf weight, SLA) for 48 species.

